# How much does habitat isolation drive forest bird morphology?

**DOI:** 10.1101/2021.02.03.429615

**Authors:** André Desrochers, Flavie Noreau

## Abstract

Rapid environmental change caused by humans has become a major concern for wildlife conservationists. But phenotypic and evolutionary responses of species to such change may often be swift enough to prevent their collapse. Several North American bird species living in boreal forests now have more pointed wings (a proxy for sustained flight efficiency), than they had a century ago. This remarkable pattern has been hypothesized as resulting from selective pressures favoring colonization of isolated habitat. Additionally, aerodynamics predict that heavier birds can achieve faster flight, a further advantage for exploring isolated habitats. We tested whether birds establishing territories in isolated areas have more pointed wings and are heavier than con-specifics found in more densely populated areas. Wing shapes of wild-caught adults from 21 passerine bird species did not generally support this prediction. However individuals with large body mass relative to their species were found more frequently in isolated habitats. Our results offer partial support for the isolation hypothesis at the landscape scale. We encourage further work at coarser, regional, scales to further examine whether wing shape and body mass evolutionarily respond to habitat isolation.

## Introduction

Whether species can adapt rapidly enough to environmental change caused by humans has become a major concern to wildlife conservationists (Gallagher et al., 2015; Hendry et al., 2008; Rice and Emery, 2003). Evolutionary processes are often thought to be vanishingly slow, but there is abundant evidence against this perception, as shown by classic studies of industrial melanism in moths (Ford, 1937), bill size in Darwin’s finches (Grant and Grant, 1989), and avian migratory behavior (Berthold et al., 1992). In fact, contemporary evolution not only occurs, but it may sometimes be key to species persistence (Bell and Gonzalez, 2009; Kinnison and Hairston, 2007; Shultz et al., 2005).

Humans have become a major, and observable, evolutionary force, by altering habitats at large scales, in particular for agriculture and forestry (Schoener, 2011). As a result, species associated with those habitats may experience changing levels of isolation from their conspecifics. Such changes in isolation may contribute to genetic structure e.g., with recruits experiencing levels of isolation similar to that of their parents because of habitat preferences modulated by ‘imprinting’ (Selonen et al., 2007; Stamps et al., 2009). Even species with high dispersal capability, such as songbirds, may exhibit reduced gene flow due to habitat preferences (Porlier et al., 2012). Thus, spatial patterns in phenotypic traits can provide insight into the factors that drive population differentiation and persistence, and may ultimately contribute to speciation (Edelaar et al., 2012).

Morphological traits offer a rich source of material to advance our understanding of processes resulting from natural and anthropogenic changes in landscapes (Tellería et al., 2013). Wings with relatively long primary feathers are associated with efficient sustained ight (Bowlin and Wikel- ski, 2008; Dawideit et al., 2009) and long migration distances (Mönkkönen, 1995). Wing shape exhibits intraspecific spatial variation among populations, migrants typically having more pointed wings than their sedentary conspecifics (Egbert and Belthoff, 2003; Palmer, 1900). There are also theoretical and empirical reasons to believe that high body mass increases flight speed through increased wing loading (Alerstam et al., 2007). Both flight efficiency and speed should therefore be considered as relevant in the morphological tradeoffs associated with dispersal and migration, even though their relative importance is unknown (Vágási et al., 2016).

Besides the importance of flight-related morphology for migration and dispersal, it is also thought to be evolutionarily labile (Piersma et al., 2005; but see Vágási et al., 2016). Striking temporal changes in wing shape occurred in forest songbirds of eastern North America over the last century (Desrochers, 2010). After ruling out the possible confounding effect of changes in migratory distances, Desrochers (2010) hypothesized that those morphological changes resulted from a selective pressure at the landscape scale, favoring greater dispersal ability in landscapes that lost significant amounts of habitat, and the converse where habitat cover significantly increased. Dispersal ability is difficult to measure, but wing shape and body mass are appropriate proxies, based on the above findings. Two key predictions resulting from Desrochers’ (2010) hypothesis is that individuals settling in landscapes with little habitat will have more pointed wings and larger body masses than conspecifics found in landscapes with abundant habitat. Such non-random spatial distribution would facilitate assortative mating by wing shape and body mass, and in turn temporal change, i.e. evolution, of this trait.

We examined whether intraspecific variation in wing shapes and body masses of forest birds of north-eastern North America is correlated with the amount of habitat at the landscape scale. More specifically, we assessed the relationship between wing pointedness, expressed as Kipp’s index (Lockwood et al., 1998), body mass, and species abundance at the landscape scale in 21 songbird species including residents and migrants, in a managed boreal forest of southern Quebec, Canada.

## Methods

We conducted field work at 30 sites separated by > 1.4 km (mean 3.0 km) at Forêt Montmorency, Québec, Canada, during the summers of 2013 and 2014 (Figure 1). This forest has been managed for timber since the 1930s, leaving a patchwork of small (10 ha) to extensive (100 ha) even-aged forest stands. Less than three percent of the surface area is composed of lakes and rivers. Young stands (< 20 years) are generally a dense mixture of deciduous and coniferous species (Mallik et al., 1997). Older stands are dominated by balsam fir (*Abies balsamea*), with occasional spruce (*Picea glauca, P. mariana*), birch (*Betula papyrifera*) and poplar (*Populus tremuloides, P. balsamifera*) groves.

**Figure 1:**
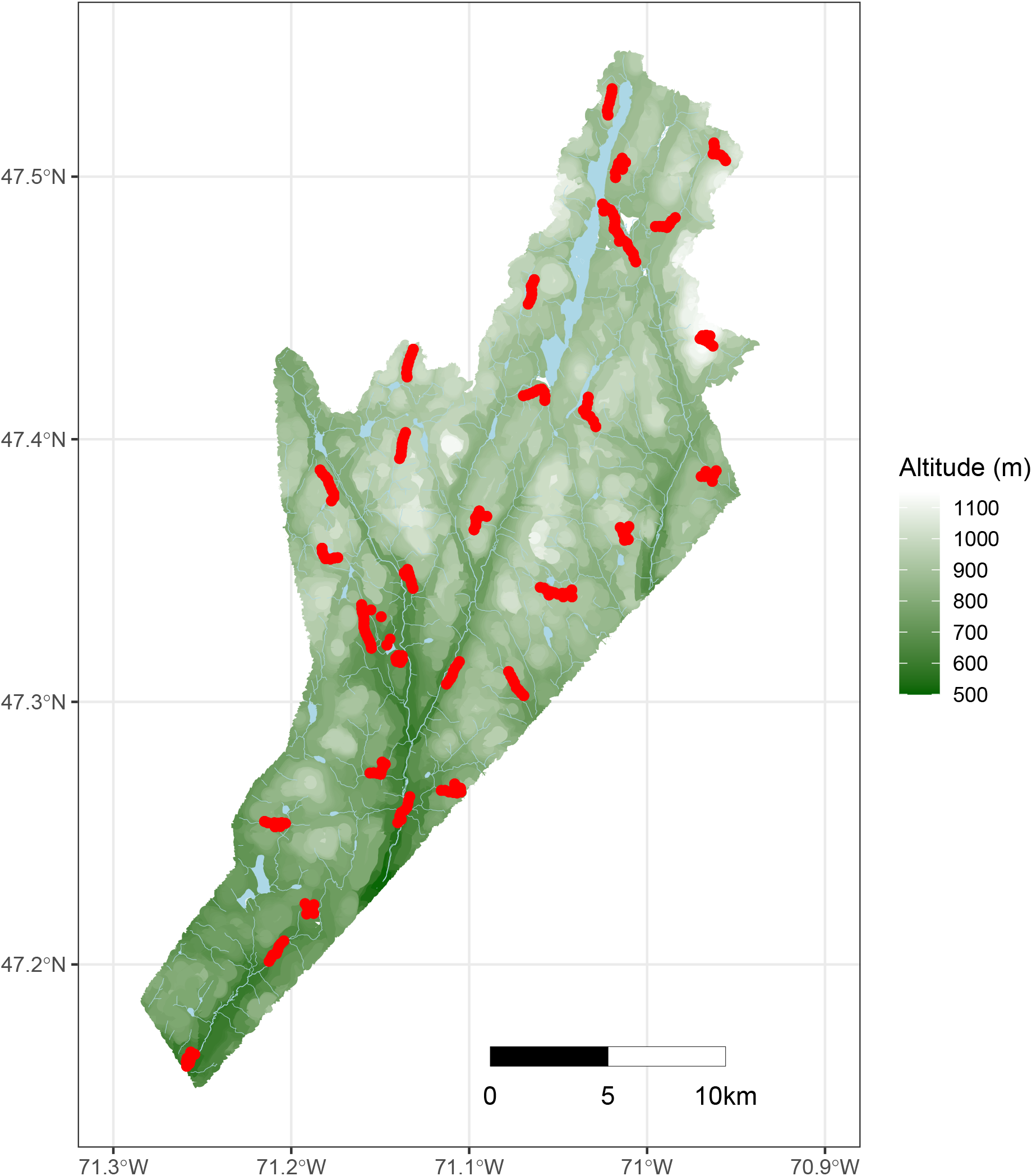
Study area. Red areas denote the location of the 30 sites where birds were counted, caught and banded.

### Field methods

At each of the 30 sites, we conducted 7 to 15 point counts with playbacks of mobbing calls to elicit responses and enhance bird detection (Corbani et al., 2014). We clustered point count stations around each site centre, with a minimum distance of 150 m from the nearest point count, and surveyed them on 2 to 3 consecutive days. We identified and added to the database all birds seen or heard within 100 m during each 15-minute point count. For each species, we retained the maximum count of individuals at each point count station, and calculated the mean count at each site. Birds caught in mist-nets (see below) were not added to birds detected in point counts. We interpreted the resulting mean counts as an index of species’ abundance at the landscape level, assuming no large differences in species detection probability among landscapes. We believe that a species’ abundance at the landscape level is a simple and effective proxy for the amount of its habitat at that scale.

At the centre of each site, we deployed mist-nets between 5:00 and 11:00 on the same dates as point counts. We used recordings of mobbing calls (Corbani et al., 2014) to increase capture rates. We banded, identified and measured birds according to a standard protocol (North American Banding Council, 2001): body mass, atten wing lentgh, fat score. We determined age and sex by plumage, the presence of a cloacal protuberance or a brood patch (Pyle, 1997). To facilitate the measurements of wing morphology, we took one to three photos of the right wing flattened on a wing ruler for each bird, with a Nikon D80 digital SLR with a resolution of 10 Megabytes in 2013 and Nikon D7100 DSLR 24 megabytes in 2014. We took three measurements in pixels from each photo with ImageJ (imagej.nih.gov): first, a 100 mm section of the wing ruler for scale, second, the total length of the wing, and third, the length between the most distal secondary feather and the front end of the wing (Lockwood et al., 1998; more details in Noreau and Desrochers, 2018).

Repeated (blind) estimates of the primary projection of the same bird by the same author had a mean standard deviation of 0.26mm (*n* = 682 birds). The mean standard deviation of primary projection estimates of the same bird measured separately by the two authors was 0.24 mm (*n* = 35 birds), corresponding to a repeatability of Kipp’s index of 99.8% (n = 35 birds). We assume that precision and accuracy of primary projection estimates were sufficiently high to yield useful intraspecific comparisons.

### Statistical analysis

To obtain a general assessment of the wing shape vs. isolation relationship, we expressed local abundance of each species at each site as standardized normal deviates (z-scores) within species. Similarly, we expressed body mass and primary projection as within-species z-scores for each individual. We preferred intraspecific z-scores in simple models over generalized least-squares and phylogenetic contrasts often found in cross-species analyses (Harvey and Pagel, 1991), because we were interested in *intra*specific, not *inters*pecific, differences in mean trait sizes. We retained species with at least 10 individuals caught, to calculate meaningful z-scores. Thus, sites with a low z-score for a given species were considered as isolated relative to sites with a higher z-score. Individuals with high z-scores for primary projection or mass were assumed to be capable of more efficient or faster sustained flight, respectively, than the average conspecific.

We considered possible bird age and sex effects on body mass (Pyle, 1997) and primary projection (la Hera et al., 2014; Mulvihill and Chandler, 1990) by repeating analyses with z-scores within age and sex groups of each species. Ages were reduced to three classes: second calendar year, ‘SY’ vs. after second calendar year, ‘ASY’, or unknown (‘AHY’). To examine interspecific differences in responses to abundance, we modelled the statistical interaction between species and local abundance effects. In addition to the above models, single-species models were run to document mprphology - abundance relationships for each species. We performed all data analyses with R (R Core Team, 2020).

## Results

We recorded 8781 birds meeting distance criteria (i.e. 100 m) at point counts, and caught and measured primary projections of 1047 individual birds (Table 1). Primary projection measurements from 1000 birds of 21 species were retained for analysis, after excluding birds of species with fewer than 10 individuals caught.

**Table 1:** Numbers of birds caught, by sex ans age class.

| Age | Female | Male | Unknown | Total |
| --- | --- | --- | --- | --- |
| After Hatch Year | 95 | 196 | 52 | 343 |
| After Second Year | 97 | 231 | 16 | 344 |
| Hatch Year | 0 | 0 | 9 | 9 |
| Second Year | 94 | 247 | 10 | 351 |
| Total | 286 | 674 | 87 | 1047 |

Primary Projection varied greatly among species, ranging from 15.9 % to 29.9 % of wing length (Table 2). The coefficient of variation of primary projection among individuals averaged 9.4 %.

**Table 2:** Primary projection, expressed as percent of wing length (*±* SD (n)). Species ordered by decreasing primary projection.

| Scientific Name | Primary Projection |
| --- | --- |
| *Catharus ustulatus* | $29.9 \pm 1.8$ (107) |
| *Setophaga striata* | $29.3 \pm 1.8$ (128) |
| *Setophaga castanea* | $28.5 \pm 1.7$ (25) |
| *Leiothlypis peregrina* | $26.7 \pm 1.6$ (17) |
| *Vireo philadelphicus* | $23.9 \pm 2.4$ (26) |
| *Setophaga virens* | $23.8 \pm 1.6$ (39) |
| *Setophaga coronata* | $23.3 \pm 1.8$ (70) |
| *Empidonax alnorum* | $22.6 \pm 1.5$ (20) |
| *Passerella iliaca* | $21.8 \pm 1.8$ (11) |
| *Empidonax flaviventris* | $21.7 \pm 2.6$ (14) |
| *Oreothlypis ruficapilla* | $21.5 \pm 2.1$ (44) |
| *Regulus satrapa* | $21.3 \pm 2.3$ (22) |
| *Junco hyemalis* | $19.7 \pm 2.3$ (51) |
| *Regulus calendula* | $19.5 \pm 2.2$ (50) |
| *Setophaga ruticilla* | $19.3 \pm 2.4$ (36) |
| *Setophaga magnolia* | $19.3 \pm 1.7$ (163) |
| *Wilsonia pusilla* | $18.9 \pm 1.8$ (29) |
| *Melospiza lincolnii* | $18.1 \pm 1.9$ (30) |
| *Zonotrichia albicollis* | $16.3 \pm 2$ (93) |
| *Geothlypis trichas* | $16.3 \pm 2.1$ (12) |
| *Poecile hudsonicus* | $15.9 \pm 2$ (13) |

Primary projection, expressed as z-scores, was similar between ages (SY vs. ASY; *β* = −0.05 *±* 0.74, *t* = −0.07, *P* = 0.94), and sexes (*β* = −0.15 *±* 0.12, *t* = −1.27, *P* = 0.21). There was no general relationship between primary projection and local abundance, both expressed as z-scores (*β* = 0.019 *±* 0.032, *t* = 0.603, *P* = 0.547; Figure 2). Adding species ID and its interaction with abundance did not improve model fit (delta deviance = 32.7, delta *df* = 40, *P* = 0.79). Nine of the 21 species had projection-to-abundance relationships consistent with the prediction (Figure 3), but none of those relationships were significant (*α* = 0.05) after adjusting for false discovery rates, respectively (Benjamini and Hochberg, 1995).

**Figure 2:**
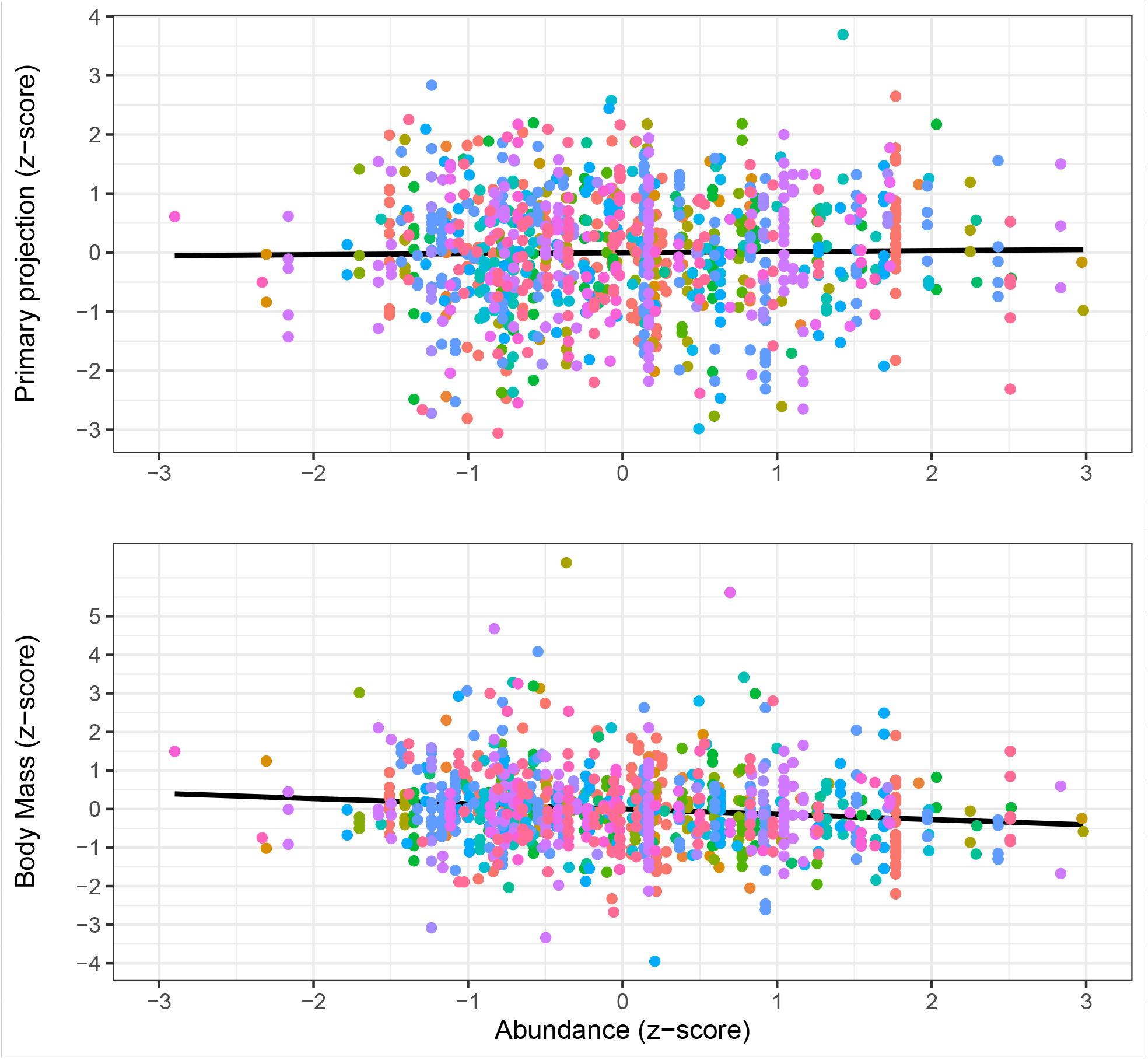
Relationships between Primary Projection, Body Mass, and landscape abundance. All variables were scaled (z-scores) to facilitate interspecific comparisons. Each of the 21 species is represented by a different colour.

**Figure 3:**
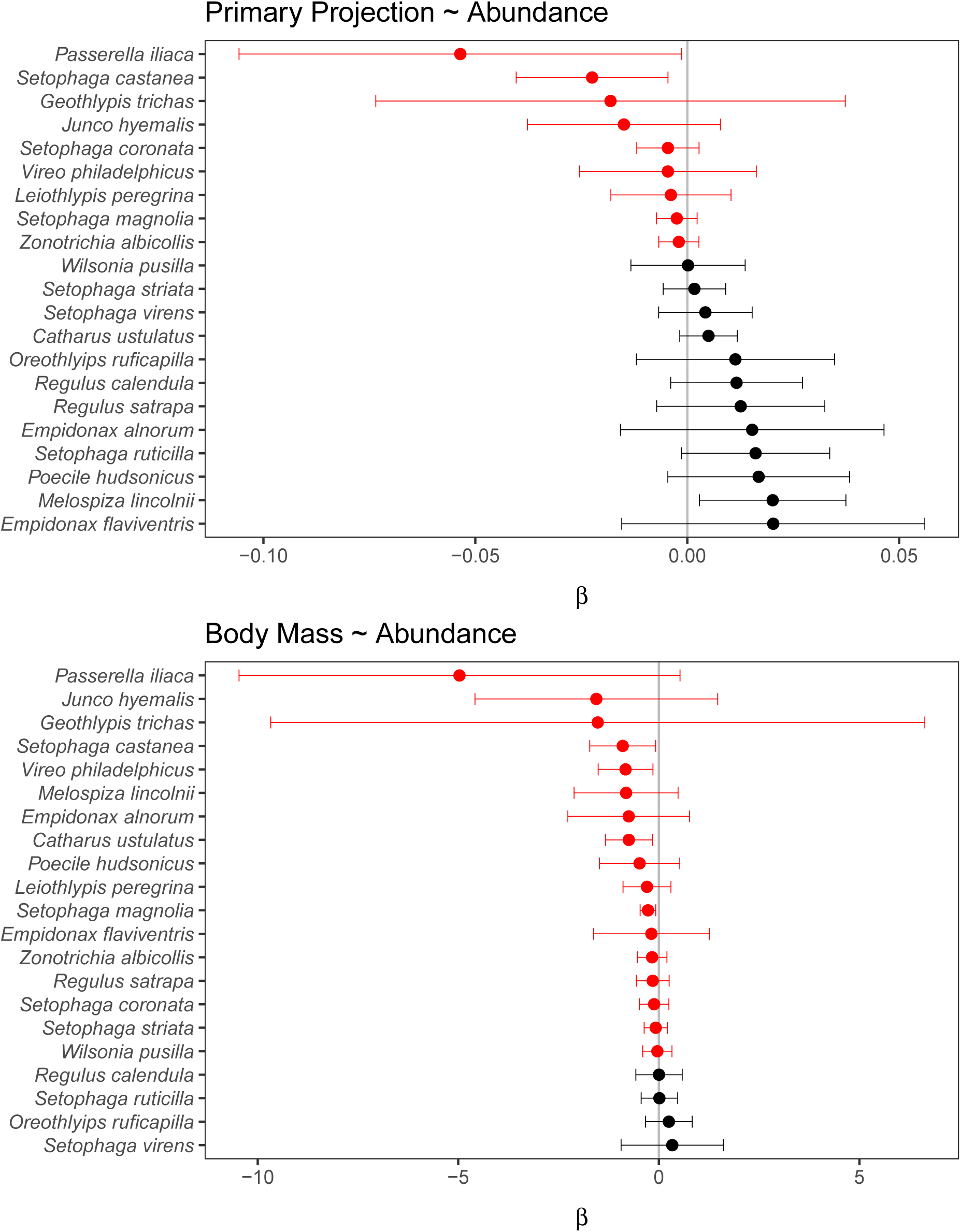
Regression estimates for the relationships between Primary projection, Body Mass, and species abundance at the landscape level. Confidence limits (95%) are shown. Negative estimates in red.

There was a general relationship between body mass and local abundance, both expressed as z-scores (*β* = −0.134 *±* 0.032, *t* = −4.168, *P* = 0; Figure 2). Adding species ID and its interaction with abundance did not improve model fit (delta deviance = 32.7, delta *df* = 40, *P* = 0.79). Seventeen of the 21 species had projection-to-abundance relationships consistent with the prediction (Figure 3), but again none of those relationships were significant (*α* = 0.05) after adjusting for false discovery rates, respectively. To avoid age and sex confounds, we repeated the analysis with z-scores for primary projection and mass within age and sex groups, and found similar results.

## Discussion

Striking temporal changes in wing shape occurred in forest songbirds of the study region over the last century (Desrochers, 2010). The latter study hypothesized that those changes resulted from a selective pressure favoring increased dispersal ability in landscapes that lost significant amounts of habitat, and the converse where habitat cover significantly increased. If true this suggests that habitat availability, inferred from species’ abundance at the landscape scale, could also explain intraspecific spatial structure in wing shape, with longer-winged individuals more likely to be found in landscapes with few conspecifics. Yet, despite substantial sample sizes, our results failed to support the wing shape hypothesis. However, for most species, heavier birds were more likely to be found in isolated habitat, although the general relationship was weak. The latter finding is consistent with the fact that birds with higher wing load are capable of faster flight (time-saving; Vágási et al. (2016)), which presumably becomes an asset when habitat selection must take place in a limited time period, among isolated habitats. It is noteworthy that Bay-breasted Warbler (*S. castanea*) was among the most responsive species with both wing shape and body mass estimates; this is the most sensitive species to habitat loss at the landscape scale in the study area (Drolet et al., 1999).

Of course, addressing a complex ecological question with a simple approach will overlook important factors and will often lead to weak relationships. It would be unrealistic to expect that phenotypic plasticity, gene flow, and reaction norms do not differ strongly among species. Furthermore, substantial intraspecific variation may occur in migration distance, which are known to induce variation in wing shape (Egbert and Belthoff, 2003), and intraspecific variation may itself vary among species. Other phenomena known to influence wing shape and body mass at the interspecific level. In the case of wing shape, resource scarcity (Kaboli et al., 2007) and foraging behavior (birds: Marchetti and Price (1995), Noreau and Desrochers (2018), bats: Norberg and Rayner (Norberg and Rayner, 1987)), are likely to occur at the intraspecific level, and may create substantial heterogeneity. Body mass is likely influenced by an even greater array of factors outside the scope of the current study. With such extraneous influences on wing shape and body mass, it is not surprising to find a large amount of statistical noise in general patterns at the level of entire species assemblages, such as thoses illustrated in Figure 2. Yet, in evolutionary terms, even weak benefits to certain morphological traits may lead to rapid evolution, given high heritabilities of most of these traits (Åkesson et al., 2008).

The weakness of the general pattern in the spatial distribution of wing shapes and body masses may also result from the scale at which the study was conducted. Besides the fact that Desrochers’ (2010) prediction was explicitly at the landscape scale, the landscape scale seemed a logical starting point for ecological reasons alone, since natal dispersal is generally thought to unfold at this spatial scale, at least in the case of songbirds (Greenwood and Harvey, 1982). However, Desrochers’ (2010) hypothesis does not apply to one particular spatial scale. For example, contemporary evolution in wing shape may be driven by bird settlement and vagrancy patterns at much coarser, regional, scales, which would explain the resilience of patterns among species documented by Desrochers (2010), in addition to well-documented regional differences within species (Peiro (2003) and references therein).

Our work could be criticized as not having examined habitat isolation per se, despite the initial thrust of the study, i.e. habitat loss. In response to this we point out that measuring habitat isolation in a cross-species analysis is difficult at best, for no two species share identical habitat. In the unlikely case that a well-defined habitat could be determined for each species, such a determination would have to be made by occupancy (Mackenzie et al., 2002) or similar models, but those models would nevertheless be based on species abundances. Thus, we believe that using a landscape-scale index of abundance is more direct than inferring it from habitat.

Contemporary evolution of traits such as wing shape (Bitton and Graham, 2015; Desrochers, 2010; Yom-Tov et al., 2006) is unsurprising, given the changing selective pressures and high heritability of morphological traits. For example, a detailed analysis of trait heritability in the Great Reed Warbler (*Acrocephalus arundinaceus*) estimated a heritability of wing length > 0.7 and primary projection > 0.4 (Åkesson et al., 2008). Given sufficient genetic variation, species can therefore respond adaptively to rapid environmental change caused by humans. This is evidenced by recent changes in migratory phenology (Miller-Rushing et al., 2008) but interestingly those changes may be inhibited by specialized wing morphology (Møller et al., 2017). The potential for rapid evolution should be recognized, lest species suffer from mismanagement (Ashley et al., 2003; Bell and Gonzalez, 2009; Rice and Emery, 2003). Unfortunately, current homogenization of landscapes by humans may thwart potential for rapid evolution, by reducing intraspecific variation, as seems to be the case in a tropical African songbird, *Andropadus virens* (Freedman et al., 2010), and possibly North-American songbirds (Martin et al., 2017).

This study provides limited support to Desrochers’ (2010) hypothesis, thus further testing for spatial patterns in wing shape and body mass remains worthwhile at similar and coarser, regional, scales. It would seem the logical next step in the study of causes and consequences of intraspecific variation in wing shape and body mass. It would contribute to a more “eco-evolutionary” approach to species management, advocated by Kinnison and Hairston (Kinnison and Hairston, 2007), and a greater recognition of mechanisms likely to drive contemporary evolution of wildlife in response to rapid environmental change.

## Acknowledgments

This research was funded by a Natural Science and Engineering Research Council of Canada’s Discovery Grant to the author. All field work was conducted under the guidelines of Université Laval’s Animal Care Committee (certificate # 2013030-2). We thank Pierre-Alexandre Dumas, Vanessa Dufresne, Jean-Michel Chabot and Martine Lapointe for help in the field. Amanda E. Martin provided helpful criticism on an earlier version of this article.

